# SbCYP79A61 Produces Phenylacetaldoxime, a Precursor of Benzyl Cyanide and Phenylacetic Acid in *Sorghum bicolor*

**DOI:** 10.1101/2022.05.11.491506

**Authors:** Veronica C. Perez, Ru Dai, Breanna Tomiczek, Jorrel Mendoza, Emily S.A. Wolf, Alexander Grenning, Wilfred Vermerris, Anna K. Block, Jeongim Kim

## Abstract

Aldoximes are amino acid derivatives that serve as intermediates for numerous specialized metabolites including cyanogenic glycosides, glucosinolates, and auxins. Aldoxime formation is mainly catalyzed by cytochrome P450 monooxygenases of the 79 family (CYP79s) that can have broad or narrow substrate specificity. Aldoxime biosynthesis in the cereal sorghum (*Sorghum bicolor*) is not well characterized. This study identified nine CYP79-encoding genes in the genome of sorghum. A phylogenetic analysis of CYP79 showed that SbCYP79A61 formed a subclade with maize ZmCYP79A61, previously characterized to be involved in aldoxime biosynthesis. Functional characterization of this sorghum enzyme using transient expression in *Nicotiana benthamiana* and stable overexpression in *Arabidopsis thaliana* revealed that SbCYP79A61 catalyzes the production of phenylacetaldoxime (PAOx) from phenylalanine, but unlike the maize enzyme, displays no detectable activity against tryptophan. Additionally, targeted metabolite analysis after stable isotope feeding assays revealed that PAOx can serve as a precursor of phenylacetic acid (PAA) in sorghum and identified benzyl cyanide as an intermediate of PAOx-derived PAA biosynthesis in both sorghum and maize. Taken together, our results demonstrate that SbCYP79A61 produces PAOx in sorghum and may serve in the biosynthesis of other nitrogen-containing phenylalanine-derived metabolites involved in mediating biotic and abiotic stresses.

**Highlight:** Identification of SbCYP79A61, an enzyme catalyzing the production of phenylacetaldoxime (PAOx) in sorghum, and identification of benzyl cyanide as an intermediate of PAOx-derived phenylacetic acid (PAA) biosynthesis in monocots.

## INTRODUCTION

Plants produce a wide variety of specialized metabolites that are structurally and functionally diverse and that play crucial roles in plant stress adaptation. Aldoximes are nitrogen-containing imines derived from amino acids that serve as precursors of various specialized metabolites including cyanogenic glycosides and glucosinolates, which function in plant defense against herbivores and pathogens (Sørensen *et al*., 2018). For example, the tyrosine-derived aldoxime 4-hydroxyphenylacetaldoxime is a precursor of dhurrin in sorghum (*Sorghum bicolor*) (Koch *et al*., 1995; Sibbesen *et al*., 1995), triglochinin in *Triglochin maritima* (Nielsen and Møller, 1999), taxiphyllin in *Taxus baccata* (Luck *et al*., 2017) and tyrosol in *Sinapis alba* (Kindl and Schiefer, 1971). The phenylalanine-derived aldoxime phenylacetaldoxime (PAOx) is a precursor of the cyanogenic glycosides amygdalin and prunasin in species such as almond (*Prunus dulcis*), apricot (*Prunus meme*) and apples (*Malus domestica*) (Yamaguchi *et al*., 2014; Bolarinwa *et al*., 2015; Hansen *et al*., 2018; Thodberg *et al*., 2018). Aliphatic aldoximes also participate in cyanogenic glycoside biosynthesis, serving as intermediates in the conversion of valine to linamarin in flax (*Linum usitatissimum*) and cassava (*Manihot esculenta*) (Tapper *et al*., 1967; Andersen *et al*., 2000) as well as in the conversion of isoleucine to lotaustralin in cassava and lima bean (*Phaseolus lunatus*) (Andersen *et al*., 2000; Lai *et al*., 2020). In Brassicales, aldoximes are precursors of glucosinolates (Underhill, 1967; Bak *et al*., 1998; Halkier and Gershenzon, 2006; Blažević *et al*., 2020) and the phytoalexin camalexin (Glawischnig *et al*., 2004; Glawischnig, 2007). Various volatiles including 2-phenylethanol (Dhandapani *et al*., 2019), (2-nitroethyl)benzene (Yamaguchi *et al*., 2021) and nitriles are generated from aldoximes and contribute to the herbivore-induced volatile blend of many species (Irmisch *et al*., 2013, 2014; McCormick *et al*., 2014; Yamaguchi *et al*., 2016; Luck *et al*., 2016).

Aldoxime production from regular or chain-elongated amino acids is catalyzed mainly by cytochrome P450 enzymes belonging to family 79 (CYP79). For example, CYP79A2 from Arabidopsis and CYP79D73 from *Plumeria rubra* catalyze PAOx production from phenylalanine (Wittstock and Halkier, 2000; Dhandapani *et al*., 2019), CYP79B2 and CYP79B3 from Arabidopsis catalyze indole-3-acetaldoxime (IAOx) production from tryptophan (Mikkelsen *et al*., 2000; Zhao *et al*., 2002), and CYP79F1 and CYP79F2 act upon chain-elongated methionine (Chen *et al*., 2003) in Arabidopsis to produce aliphatic aldoximes that are precursors of aliphatic glucosinolates (Harun *et al*., 2020). Other CYP79 enzymes have greater promiscuity, such as poplar (*Populus trichocarpa*) CYP79D6 and CYP79D7, which can generate aldoximes from tryptophan, phenylalanine, isoleucine, leucine, and tyrosine (Irmisch *et al*., 2013). Recently, heterologous expression analyses demonstrated that the Arabidopsis enzymes CYP79C1 and CYP79C2 have promiscuous aldoxime production activity, with CYP79C2 acting mainly upon leucine and phenylalanine, but with minor activity towards isoleucine, tryptophan and tyrosine, whereas CYP79C1 acts upon leucine, isoleucine, phenylalanine and valine (Wang *et al*., 2020). CYP79 enzymes are widely distributed in both monocots and dicots and have been identified in some gymnosperms (Sibbesen *et al*., 1995; Nielsen and Møller, 2000; Irmisch *et al*., 2013, 2015; Yamaguchi *et al*., 2014, 2021; Luck *et al*., 2016, 2017; Hansen *et al*., 2018; Sørensen *et al*., 2018; Thodberg *et al*., 2018; Dhandapani *et al*., 2019; Lai *et al*., 2020). There is also evidence for other flavin-containing monooxygenases (FMO) that are capable of catalyzing aldoxime formation. Examples of such FMOs have been identified in fern and shown to be involved in PAOx formation (Thodberg *et al*., 2020).

Aldoximes also play crucial roles in plant growth. Specifically, IAOx and PAOx are precursors of the two major auxins indole-3-acetic acid (IAA) and phenylacetic acid (PAA), respectively, in Brassicales and maize (*Zea mays*), although the molecular components of aldoxime-derived auxin biosynthesis remain to be elucidated (Zhao *et al*., 2002; Sugawara *et al*., 2009; Aoi *et al*., 2020; Perez *et al*., 2021).

In maize, aldoxime-derived auxin biosynthesis is initiated by ZmCYP79A61, which is capable of catalyzing both IAOx and PAOx formation (Irmisch *et al*., 2015). Sorghum (*Sorghum bicolor*) is a monocot that is closely related to maize (Schnable *et al*., 2009) and cultivated globally for feed, food, fodder, and biofuel production (Paterson *et al*., 2009). Sorghum is known to contain one aromatic aldoxime production enzyme SbCYP79A1, which acts upon tyrosine and is involved in the production of the defense compound dhurrin (Sibbesen *et al*., 1995). Given the role of aldoximes in plant responses to biotic and abiotic stresses, and the growing interest in sorghum as a climate-resilient crop, a better understanding of aldoxime metabolism in sorghum will enable novel crop improvement strategies.

In this study we mined the sorghum genome for *CYP79* homologs and identified nine candidate sequences, including one homolog closely related to the maize gene encoding ZmCYP79A61. Further characterization of this candidate (*SbCYP79A61*) via genetic studies and transient expression identified it as an enzyme capable of producing PAOx. Additionally, our stable isotope labelling assays identified that sorghum produces PAA from PAOx and that benzyl cyanide is an intermediate of PAOx-derived PAA production in both sorghum and maize.

## MATERIALS and METHODS

### Plant material and growth conditions

*Arabidopsis thaliana* Col-0, maize B104, sorghum RTx430 and *Nicotiana benthamiana* were used as wild-type plants. After 3 days of cold treatment at 4°C, Arabidopsis seeds were planted on soil and kept in a growth chamber at 21°C with 16h light/8h dark. Sorghum and *N. benthamiana* seeds were planted directly on soil and kept at 25°C with 16h light/8h dark. Maize seeds were imbibed with water for 1h before being planted on soil and kept at 25°C with 16h light/8h dark. Arabidopsis double mutant *cyp79b2 cyp79b3* (*b2b3*) and its genotyping method are described in Zhao *et al*. (2002).

### CYP79 sequence analysis and phylogenetic tree generation

BLASTp analysis (Altschul *et al*., 1990) was performed with the *Sorghum bicolor* v3.1.1 genome from Phytozome v12 (phytozome.jgi.doe.gov) using the protein sequence of maize ZmCYP79A61 as the query. Protein sequences of sorghum, maize, poplar, plumeria and Arabidopsis CYP79 proteins were aligned using the MUSCLE algorithm (Edgar, 2004) (gap open, -2.9; gap extend, 0; hydrophobicity multiplier, 1.2; clustering method, UPGMA) implemented in Mega7 (Kumar *et al*., 2018). Phylogenetic trees were constructed with Mega7 using a neighbor-joining algorithm (model/method, Poisson; substitution type, amino acid; rates among sites, uniform; gaps/missing data treatment, pairwise deletion deletion). A bootstrap resampling analysis with 1000 replicates was performed to evaluate tree topology.

### Plasmid construction and transgenic plant growth

To generate *SbCYP79A61* overexpression constructs, the *SbCYP79A61* open reading frame was synthesized within the pUC57 vector (Genewiz, South Plainfield, NJ). The synthesized entry vector was subsequently recombined with the destination vector pCC0995 (Weng *et al*., 2010), in which expression of the open reading frame is under the control of the cauliflower mosaic virus (CaMV) 35S promoter to generate the *35S::SbCYP79A61* construct. The *35S::SbCYP79A61, 35S::ZmCYP79A61* (Perez *et al*., 2021), and *35S::AtCYP79B2* (Zhang *et al*., 2020) constructs were introduced into *Agrobacterium tumefaciens* (GV3101) following a method described by Zhang *et al*. (2020). All three constructs were used to transiently express CYP79 enzymes in *N. benthamiana*, described in the following section.

The *35S::SbCYP79A61*-containing *Agrobacterium tumefaciens* GV3101 cells were additionally used to transform Arabidopsis wild type (Col-0) or *b2b3* mutants via a floral dipping method (Zhang *et al*., 2020). More than ten T1 plants were screened by application of 0.2% Basta (Rely 280, BASF, Iselin, NJ). Several single-insertion homozygous T3 lines were established based on Basta resistance.

### Transient expression in *Nicotiana benthamiana*

Suspensions of *A. tumefaciens* cells containing the *35S::SbCYP79A61, 35S::ZmCYP79A61, 35S::AtCYP79B2* or pCC0995 (vector control) constructs were infiltrated into leaves of *N. benthamiana* as previously described (Norkunas *et al*., 2018). Briefly, suspensions of transformed bacteria were adjusted to an OD_600_ of 0.6 in infiltration buffer (10mM MgCl_2_, 10mM MES, pH 5.6), and acetosyringone was added to a final concentration of 150μM. Bacterial suspensions were then incubated at room temperature for 1h and loaded into a blunt-tipped plastic syringe that was then used to infiltrate the underside of leaves of three-week-old plantlets. The three to four fully expanded leaves of independent plantlets were infiltrated with cell suspensions containing the different vectors. After three days, leaf tissue samples were harvested, extracted, and analyzed for aldoxime content.

### RNA extraction and qRT-PCR analysis

Total RNA was isolated from 2-week-old leaves by using QIAGEN RNeasy Plant Mini Kit (QIAGEN, Germantown, MD, 74904). Five μg of total RNA was treated with DNase following the manufacturer’s protocol, and 1μg RNA was used for first-strand cDNA synthesis using reverse transcription kit (ThermoFisher Scientific, MA, 4368814). qPCR reactions were performed in a 10μL volume mix using SYBR Green components (ThermoFisher Scientific, MA, A25742) in StepOnePlus Real-time PCR system (Applied Biosystems). *SbCYP79A61* transcripts were amplified with the following primers: forward primer (5′-ACCCTCTTCGCTTCAACCC-3′) and reverse primer (5′-ATGATGCTCATAGCAGTGCCG-3′). Gene expression was normalized to the reference gene *TUB3* (At5g62700), which was amplified with the following primers: forward primer (5′-TGGTGGAGCCTTACAACGCTACTT-3′) and reverse primer (5′-TTCACAGCAAGCTTACGGAGGTCA-3′). Relative expression was calculated based on the 2^-ΔΔCt^ method (Livak and Schmittgen, 2001). qRT-PCR analysis was conducted with three biological replicates.

### Metabolite analysis

Glucosinolates were extracted from three-week-old Arabidopsis leaves following the method described in Perez *et al*. (2021). Ten μL of extracts was analyzed on an UltiMate 3000 HPLC system (ThermoFisher Scientific, MA) equipped with a diode array detector (DAD) in the UV-vis region 200-500nm. The compounds were separated on an Acclaim™ RSLC120 C18 column (100mm×3mm; 2.2μm) with a flow rate of 0.5 ml min^-1^ (ThermoFisher Scientific, MA). Solvent A (0.1% formic acid (v/v) in water) and solvent B (acetonitrile) were used as mobile phases with a gradient program as follow: 0.3-5min 5-14%, 5-8.25min 14-14.5%, 8.25-11.5min 14.5-18%, and 11.5-15min 18-95% of solvent B increase. The column temperature was 40°C. The contents of benzyl glucosinolate and indole glucosinolate were quantified based on the peak area at 220nm and the authentic standards benzyl glucosinolate (EMDMillipore Sigma, PHL89216) and indol-3-ylmethyl glucosinolate (EMDMillipore Sigma, PHL80593) were used for quantification. Soluble metabolite analyses were conducted with four biological replicates.

### Feeding assay

Leaf segments (approximately 10-12 cm) of the first leaves of 8-day-old maize and sorghum seedlings were placed in aqueous solutions containing 0.005% v/v Triton X-100 alone, with 30 μM of unlabeled or stable isotope, deuterium-labeled PAOx (containing deuterium atoms attached to benzyl ring carbons), or with 30μM of labeled or unlabeled benzyl cyanide. Unlabeled benzyl cyanide and labeled benzyl cyanide (benzyl cyanide-α-^13^C) were purchased from EMDMillipore Sigma (B19401 and 486973). Labeled and unlabeled PAOx were synthesized as described in Perez *et al*. (2021). Feeding assays were conducted with three to four biological replicates. After 24 h of incubation, samples were removed from the solutions, dried, weighed, flash-frozen in liquid nitrogen and stored at -80°C until extraction.

### Aldoxime and D_5_-PAA detection using LC-MS

Aldoxime synthesis, aldoxime and D_5_-PAA extraction, and D_5_-PAA detection were performed using methods described in Perez *et al*. (2021). Briefly, 50-100mg fresh weight samples were extracted in 1mL cold sodium phosphate buffer (50mM, pH 7.0) containing 0.1% (w/v) diethyl dithiocarbamic acid sodium salt. Liquid chromatography solutions and gradient program, mass spectrometer and ionization parameters, and MRM monitoring for D_5_-PAA detection were described previously (Perez *et al*., 2021).

For IAOx and PAOx detection, all samples were resuspended in water after extraction and analyzed using Vanquish Horizon ultra-high performance liquid chromatography (UHPLC) installed with an Eclipse Plus C18 column (2.1 × 50mm, 1.8 μm) (Aligent) and mass analysis was performed using a TSQ Altis Triple Quadruple (Thermo Scientific) MS/MS system with an ion funnel. The mass spectrometer was operated in positive ionization mode at an ion spray voltage of 4800V. Formic acid (0.1%) in water and 100% acetonitrile were used as mobile phases A and B, respectively, with a gradient program (0-95% mobile phase B over 4 min) at a flow rate of 0.4 ml/min. The sheath gas, aux gas, and sweep gas were set at 50, 9, 1 (arb unit), respectively. Ion transfer tube and vaporizer temperatures were set at 325°C and 350°C, respectively. For MRM monitoring, both Q1 and Q3 resolutions were set at 0.7 FWHM with argon at 1.5 mTorr as collision-induced dissociation gas. The scan cycle time was 0.8 s. MRM was used to monitor parent ion→ product ion transitions for each analyte as follows: m/z 175.087→ 158 (CE, 16V) for IAOx and m/z 136→ 119 (CE, 14V) for PAOx.

### Quantification of PAA and benzyl cyanide and detection of labeled PAA and benzyl cyanide using GC-MS

To quantify PAA and benzyl cyanide we used a method adapted from Schmelz *et al*. (2004). Briefly, tissues were flash frozen in liquid nitrogen and 100mg were extracted in 300μl of H_2_O:1-propanol:HCl (1:2:0.005) spiked with 100ng of D_7_-PAA as an internal standard. Samples were homogenized using 1g of ceramic beads in a Fast Prep homogenizer and further extracted with 1ml methylene chloride. The samples were centrifuged and the methylene chloride:1-propanol layer was collected and derivatized with trimethylsilyldiazomethane, quenched with acetic acid in hexane and volatile compounds were collected by vapor phase extraction on SuperQ columns. Volatiles were eluted in methylene chloride and analyzed using GC-MS (GC: Agilent 7890B, MS: Agilent 5977B, column: Agilent DB-35MS [length: 30m, diameter: 0.25mm, film thickness: 0.25μm]) with chemical ionization using iso-butane. The ions monitored and retention times for the compounds are as follows: derivatized PAA (methyl phenylacetate) (m/z 151, RT 6.74 min), derivatized D_7_-PAA (methyl D_7_-phenylacetate) (m/z 158, RT 6.71 min), derivatized PAA-α-^13^C (methyl phenylacetate-α-^13^C) (m/z 152, RT 6.74 min), benzyl cyanide (m/z 118, RT 6.79 min), benzyl cyanide-α-^13^C (m/z 119, RT 6.79 min) and D_5_-benzyl cyanide (m/z 123, RT 6.78 min). Compounds were confirmed and quantified using authentic standards. Volatile compound analysis was conducted with three to four biological replicates.

### Data Analysis

Statistical analyses for all experiments were conducted in R (version 4.0.2) and RStudio (version 1.3.1056). For data sets with two categories or treatments, statistical significance was determined using Student’s t-test. For data sets with three or more categories, statistical significance was determined using analysis of variance (ANOVA) and Tukey’s test.

### Accession Numbers

Sequence data for proteins and genes discussed in this study can be found in GenBank under the following identifiers: SbCYP79A61 (*Sobic*.*010G172200*, XP_002438593.1); SbCYP79A1 (*Sobic*.*001G012300*, XP_002466099.1); SbCYP79A89 (*Sobic*.*001G185900*, OQU91462.1); SbCYP79A91 (*Sobic*.*001G187400*, XP_002464329.1); SbCYP79A92 (*Sobic*.*001G187500*, XP_002464333.1); SbCYP79A93 (*Sobic*.*001G187600*, XP_002464334.2); SbCYP79A94 (*Sobic005G157600*, OQU83670.1); SbCYP79A95 (*Sobic*.*005G158200*, XP_002449720.2); SbCYP79A96 (*Sobic*.*007G090457*, XP_002446424.1); SbCYP71E1 (*Sobic*.*001G012200*, XP_002466097.1); ZmCYP79A61 (*GRMZM2G138248*, AKJ87843.1); GRMZM2G011156 (*GRMZM2G011156*, XP_008644689.1; GRMZM2G105185 (*GRMZM2G105185*, XP_008665466.1); GRMZM2G178351 (*GRMZM2G178351*, XP_008644663.1); AtCYP79B2 (*At4g39950*, NP_001328605.1); AtCYP79B3 (*At2g22330*, NP_001323954.1); AtCYP79A2 (*At5g05260*, NP_001318481.1); AtCYP79F1 (*At1g16410*, NP_563996.2); AtCYP79F2 (*At1g16400*, NP_563995.2); AtCYP79C1 (*At1g79370*, NP_178055.2); AtCYP79C2 (*At1g58260*, NP_176122.2); AtCYP83A1 (*At4g13770*, NP_193113.1); AtCYP83B1 (*At4g31500*, NP_194878.1); PrCYP79D73 (*PrCYP79D73*, AZT89153.1); PtCYP79D6 (*Potri*.*013G157200*, AHF20912.1); PtCYP79D7 (*Potri*.*013G157300*, AHF20913.1).

## RESULTS

### Identification of CYP79 homologs in *Sorghum bicolor*

CYP79 monooxygenases catalyze aldoxime formation in many plant species. The maize genome contains four genes encoding CYP79 enzymes, one of which, ZmCYP79A61, has been shown to produce IAOx and PAOx (Irmisch *et al*., 2015). In maize, the aldoximes IAOx and PAOx act as precursors of the auxins IAA and PAA, respectively (Perez *et al*., 2021). In sorghum, however, the only CYP79 enzyme with known function is SbCYP79A1, which produces *p*-hydroxyphenylacetaldoxime, the precursor for dhurrin, from tyrosine (Koch *et al*., 1995; Sibbesen *et al*., 1995; Bak *et al*., 2000). Given the universal occurrence of auxins and the similarities in genome sequence and organization between maize and sorghum, we hypothesized that sorghum ZmCYP79A61 homologs can generate IAOx and PAOx.

To identify putative CYP79-encoding genes in sorghum, we screened the genome of *Sorghum bicolor* v3.1.1 using the Phytozome (v12) database with maize ZmCYP79A61 protein sequence as a query. This initial search identified nine candidate sequences that included SbCYP79A1 (*Sobic*.*001G012300*). According to the CYP450 nomenclature committee (Nelson, 2009), the remaining eight sequences are annotated as SbCYP79A61 (*Sobic*.*010G172200*), SbCYP79A89 (*Sobic*.*001G185900*), SbCYP79A91 (*Sobic*.*001G187400*), SbCYP79A92 (*Sobic*.*001G187500*), SbCYP79A93 (*Sobic*.*001G187600*), SbCYP79A94 (*Sobic005G157600*), SbCYP79A95 (*Sobic*.*005G158200*), and SbCYP79A96 (*Sobic*.*007G090457*).

Phylogenetic analysis of the sorghum and maize CYP79 enzymes was performed using these protein sequences, as well as characterized CYP79 enzymes from Arabidopsis, *Plumeria rubra* (Dhandapani *et al*., 2019) and poplar (*Populus trichocarpa*) (Irmisch *et al*., 2013). In the resulting phylogenetic tree, the maize and sorghum CYP79s were split among two major sub-clades (**Fig 1**). One sub-clade includes three maize CYP79s (GRMZM2G011156, GRMZM2G105185, GRMZM2G178351) and seven sorghum CYP79s (SbCYP79A89, 91 to 96). The other sub-clade includes SbCYP79A61, SbCYP79A1, and ZmCYP79A61, and is close to CYP79s from Arabidopsis, plumeria and poplar known to be involved in aromatic aldoxime production (**Fig 1**). It was previously shown that *GRMZM2G011156, GRMZM2G105185*, and *GRMZM2G178351* are not expressed in most of organs (Irmisch *et al*., 2015). Similarly, the six *SbCYP79*s in this sub-clade except for *SbCYP79A95* have low or no expression in sorghum tissues (**Fig S2**) (EMBL-EBI Expression Atlas, https://www.ebi.ac.uk/gxa/experiments/E-MTAB-3839/Result). *SbCYP79A1* is highly expressed in most tissues except flowers, whereas *SbCYP79A61* is expressed in aerial tissues including flowers and shoots (**Fig S2**). The open reading frame of *SbCYP79A61* encodes a protein of 547 amino acids, which, when aligned with maize ZmCYP79A61, revealed 84.3% amino acid identity (**Fig S1, Table S1**). The SbCYP79A61 contains motifs conserved among other CYP79A proteins such as the heme binding site ([S/T]F[S/T]TGRRGCxA), the FxP[E/D]RH motif and the NP motif (Bak *et al*., 2006) (**Fig S1**). Given the results of the expression pattern analysis and phylogenetic analysis, SbCYP79A61 is the most likely candidate for a sorghum enzyme involved in the biosynthesis of IAOx or PAOx.

**Figure 1.**
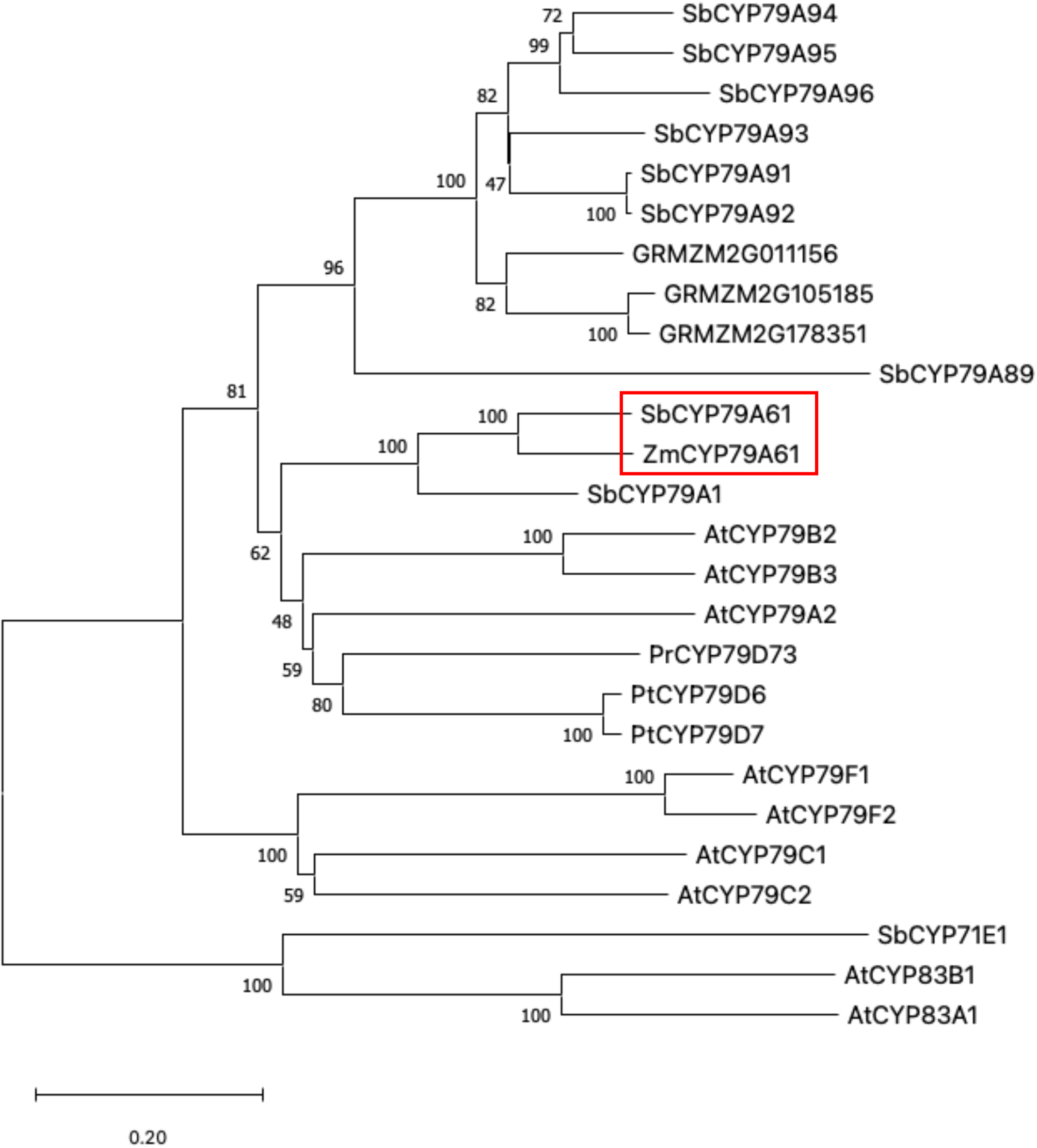
Phylogenetic tree of CYP79 sequences from sorghum, maize, Arabidopsis, *Plumeria rubra* and poplar. The red box highlights the position of the ZmCYP79A61/SbCYP79A61 subclade. The rooted tree was inferred with the neighbor-joining method and n=1000 replicates for bootstrapping. Bootstrap values are shown next to each node. As outgroups, the enzymes CYP71E1 from sorghum and CYP83A1/B1 from Arabidopsis were chosen. Accession numbers are provided in the Methods section.

### Biochemical characterization of SbCYP79A61

To determine if SbCYP79A61 can produce PAOx or IAOx, we transiently expressed *SbCYP79A61* under the control of the *CaMV35S* promoter in *Nicotiana benthamiana* (**Fig 2**). As controls for aldoxime production, *N. benthamiana* plants were infiltrated with *A. tumefaciens* harboring vectors containing *ZmCYP79A61* (for IAOx and PAOx production), *AtCYP79B2* (for IAOx production) or the vector alone as a negative control. The maize ZmCYP79A61 can act upon both tryptophan and phenylalanine, albeit with lower affinity towards tryptophan, whereas Arabidopsis AtCYP79B2 only produces IAOx (**Fig 2**) (Zhao *et al*., 2002; Irmisch *et al*., 2015). As expected, both IAOx and PAOx were observed in leaves expressing *ZmCYP79A61* and only IAOx was observed in leaves expressing *AtCYP79B2*. Interestingly only PAOx but not IAOx was detectible in leaves expressing *SbCYP79A61* (**Fig 2**), suggesting that SbCYP79A61 has activity towards phenylalanine *in vivo*.

**Figure 2.**
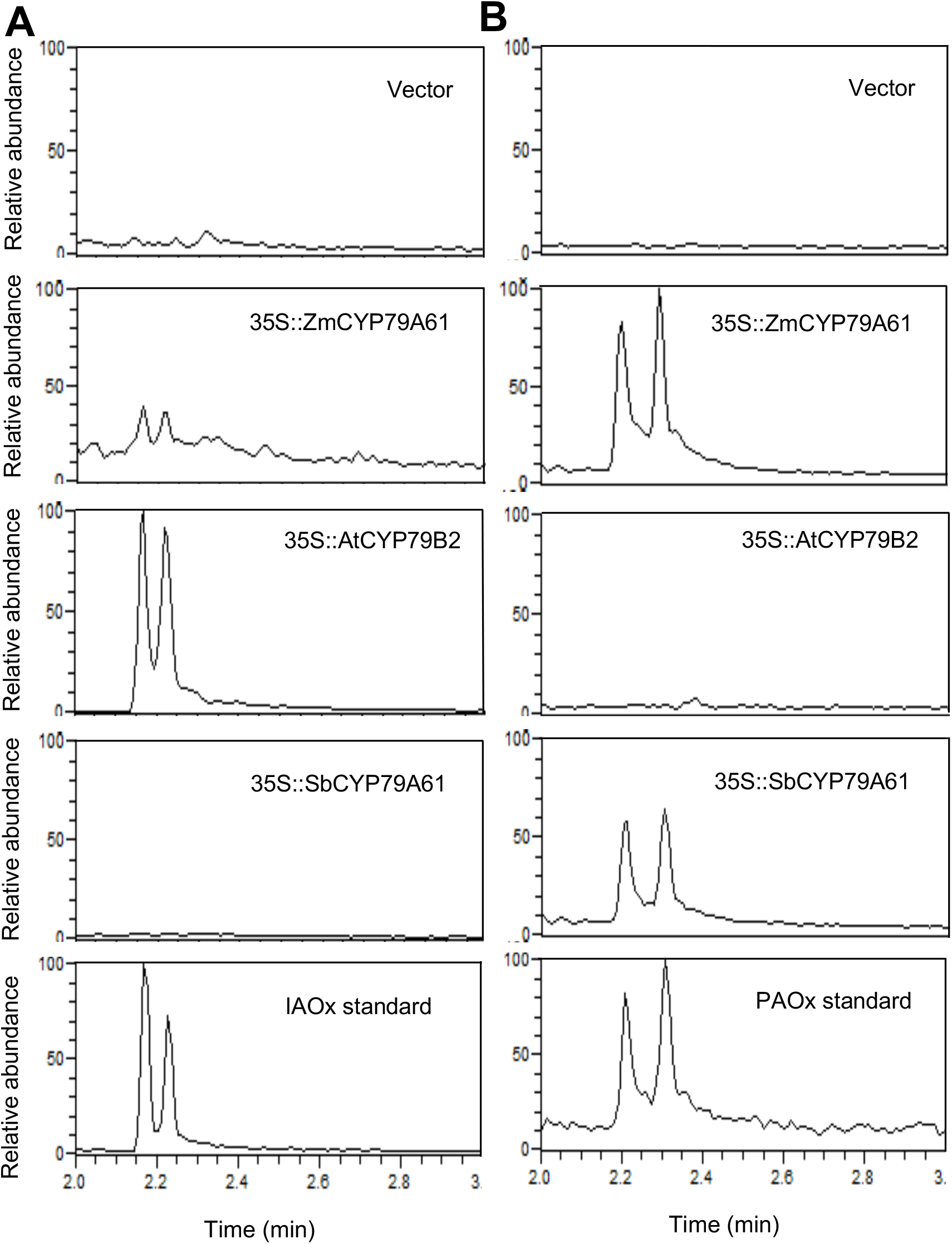
Transient expression of *SbCYP79A61* in *N. benthamiana* produces PAOx. LC-MS chromatograms for IAOx (A) and PAOx (B) in N. benthamiana leaves of 3-week-old plantlets after infiltration with Agrobacterium tumefaciens strains harboring constructs containing the CYP79 genes *ZmCYP79A61* (second from top), *AtCYP79B2* (third from top) or *SbCYP79A61* (fourth from top). Chromatograms at the top represent plants infiltrated with A. tumefaciens harboring the empty vector, and chromatograms on the bottom display aldoxime standards. Samples were taken 3 days after infiltration.

To confirm the specificity of SbCYP79A61 for phenylalanine, we generated stable Arabidopsis transformants expressing *SbCYP79A61* (**Fig 3A**) in wild type and an IAOx-deficient mutant. The Arabidopsis ecotype used in this study, Columbia-0 (Col-0), accumulates indole glucosinolates, which are derived from IAOx, but not PAOx-derived benzyl glucosinolate in leaf tissue (**Fig 3B, C**) (Kliebenstein *et al*., 2001; Perez *et al*., 2021). If SbCYP79A61 activity results in PAOx production, we hypothesized the Arabidopsis wild-type plants expressing *SbCYP79A61* to produce benzyl glucosinolates. The *cyp79b2 cyp79b3* double mutant (*b2b3*) is incapable of generating IAOx due to disruption of IAOx-producing enzymes CYP79B2 and CYP79B3, and, therefore, does not produce indole glucosinolates (**Fig 3C**) (Zhao *et al*., 2002). If, on the other hand, SbCYP79A61 activity results in IAOx production, we hypothesized the Arabidopsis *b2b3* mutants expressing *SbCYP79A61* would produce indole glucosinolates. Two overexpression lines in the wild-type background and two in the *b2b3* background were established and named *ox-1/WT, ox-2/WT, ox-11/b2b3*, and *ox-12/b2b3*. All *SbCYP79A61* overexpression lines were found to produce benzyl glucosinolate, while the wild type and *b2b3* mutant did not produce it (**Fig 3B**). This suggests that SbCYP79A61 can produce PAOx from phenylalanine. However, indole glucosinolates in *ox-11/b2b3* and *ox-12/b2b3* were below the detection limit and were not increased in lines *ox-1/WT* and *ox-2/WT* (**Fig 3C**). These results are consistent with our *N. benthamiana* infiltration results (**Fig 2**) and suggest that SbCYP79A61 has high activity towards phenylalanine, but little or no activity towards tryptophan.

**Figure 3.**
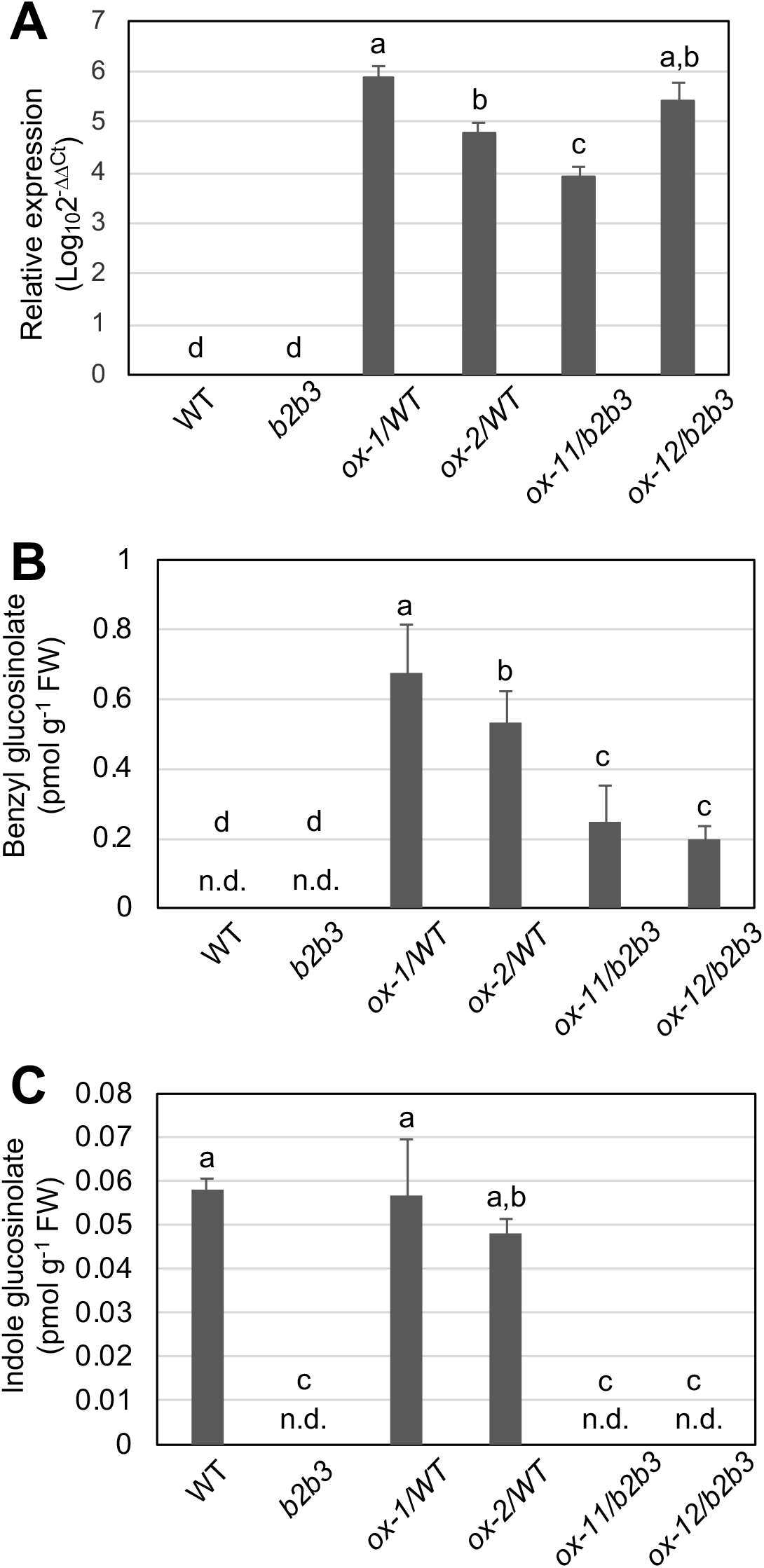
Overexpression of *SbCYP79A61* in Arabidopsis results in accumulation of benzyl glucosinolate but not indole glucosinolate. (A) Relative expression (log_10_2^-ΔΔCt^) of SbCYP79A61 in wild type, *b2b3*, and *SbCYP79A61* overexpression lines in the wild-type (*ox-1/WT, ox-2/WT*) and *b2b3* (*ox-11/b2b3, ox-12/b2b3*) genetic backgrounds (n=3). (B) Benzyl glucosinolate concentration of wild type, *b2b3*, and *SbCYP79A61* overexpression lines (n=4). (C) Indole glucosinolate (I3M) concentration of wild type, *b2b3*, and *SbCYP79A61* overexpression lines (n=4). Metabolite content was analyzed from 2-week-old whole aerial parts. Bars represent mean, and error bars represent standard deviation. The means were compared by one-way ANOVA and statistically significant differences among sample means (P < 0.05) were identified by Tukey’s test and indicated by different lowercase letters.

### Benzyl cyanide is an intermediate of the PAOx-derived PAA biosynthesis in monocots

As PAOx can act as a precursor to PAA in Arabidopsis and maize, we used feeding and isotope labeling studies to investigate the existence of this pathway in sorghum (Perez *et al*., 2021). The feeding of sorghum leaves with PAOx led to significantly increased levels of PAA compared with water-fed samples (**Fig 4A**). Additionally, when sorghum leaves were fed with D_5_-PAOx, they were found to produce D_5_-PAA (**Fig 4B-D**) upon LC-MS analysis. On the other hand, water-fed sorghum leaves were unable to produce D_5_-PAA (**Fig 4B-D**). These results suggest that sorghum, like maize and Arabidopsis, can utilize PAOx generated from SbCYP79A61 to produce PAA.

**Figure 4.**
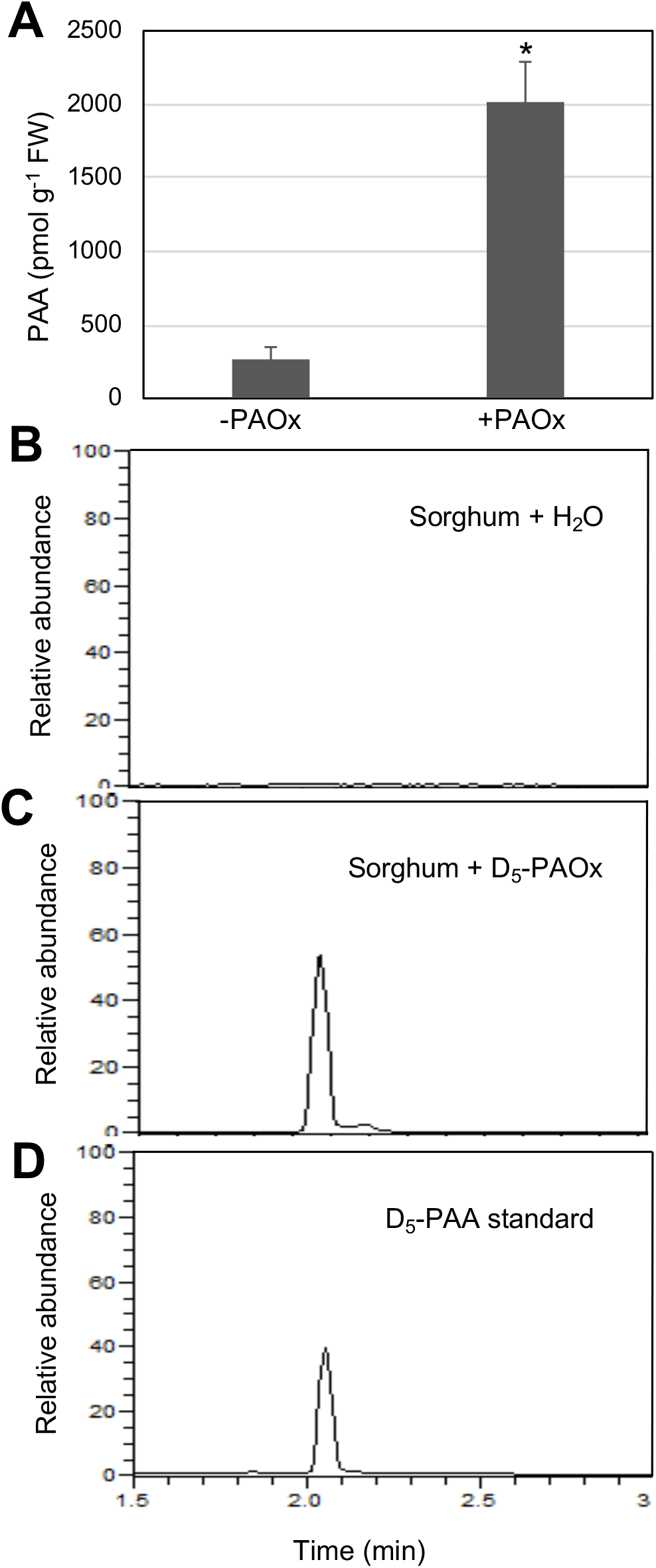
Sorghum can synthesize PAA from PAOx. (A) Free PAA content of sorghum leaves fed with water or PAOx (n=3). Bars represent mean, and error bars represent standard deviation. The means were compared by Student’s t-test and statistically significant differences (P < 0.05) are indicated by asterisks (*). (B-D) LC-MS chromatograms for the D_5_-PAA standard (D) and endogenous D_5_-PAA in sorghum leaves after feeding with water (B) or D_5_-PAOx (C). Leaf segments of 8-day-old sorghum plants were incubated with either water or 30 μM of aldoximes for 24 h.

As glucosinolate degradation produces nitriles such as benzyl cyanide and indole-3-acetonitrile, and nitrilases convert nitriles to acetic acid, it has been proposed that benzyl cyanide can be a precursor of PAA in Arabidopsis (Urbancsok *et al*., 2018). Several studies have shown that PAOx serves as a precursor of benzyl cyanide as part of the volatile emission of various species (Irmisch *et al*., 2013, 2014, 2015; Dhandapani *et al*., 2019). Thus, we hypothesized that benzyl cyanide might be derived from PAOx and an intermediate of the PAOx-derived PAA biosynthesis in sorghum. To test this hypothesis, we examined benzyl cyanide production in sorghum plants fed with PAOx. We detected a compound with a mass-to-charge ratio (m/z) of 118 that accumulated in PAOx-fed sorghum plants but was absent in water-fed plants. MS analysis identified this compound as benzyl cyanide (**Fig 5A**), which was confirmed with an authentic standard. To determine if benzyl cyanide is directly made from PAOx in sorghum, we analyzed D_5_-PAOx-fed samples for the presence of deuterium-labeled D_5_-benzyl cyanide. We detected a compound with m/z 123 and retention time matching that of benzyl cyanide in sorghum upon feeding with D_5_-PAOx, but not water (**Fig 5C-D**). As D_5_-benzyl cyanide has a mass-to-charge ratio of 123 (**Fig 5B**), these results suggest that sorghum can directly convert PAOx to benzyl cyanide.

**Figure 5.**
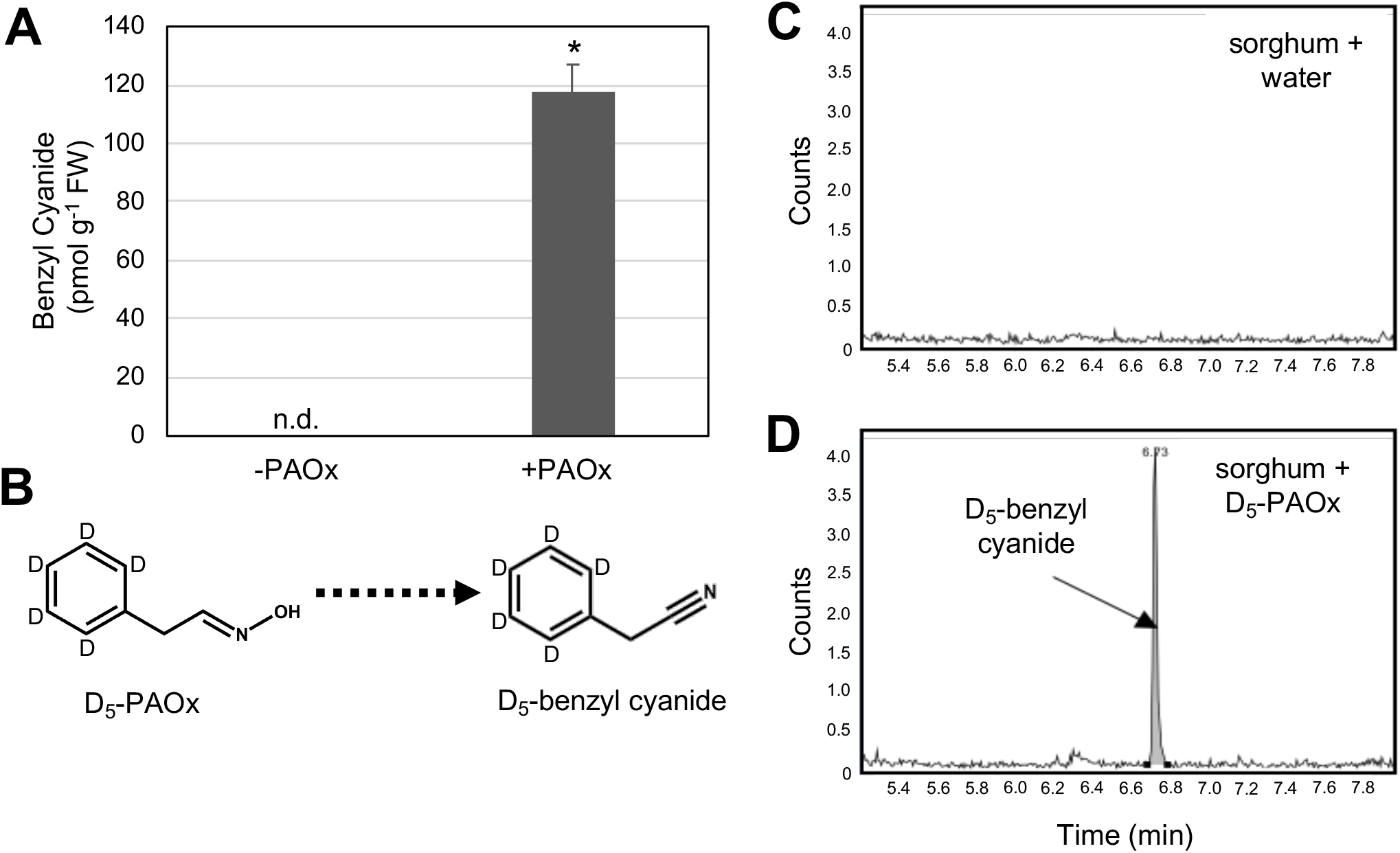
Benzyl cyanide is derived from PAOx in sorghum. (A) Benzyl cyanide concentration of sorghum leaves fed with water or PAOx (n=3). Bars represent mean, and error bars represent standard deviation. The means were compared by Student’s t-test and statistically significant differences (P < 0.05) are indicated by an asterisk (*). (B) Schematic of proposed conversion of D_5_-PAOx to D_5_-benzyl cyanide. Position of deuterium atoms in D_5_-PAOx and D_5_-benzyl cyanide are denoted by the letter ‘D’. (C-D) GC-MS chromatograms set for detection of D_5_-benzyl cyanide (m/z 123) in sorghum leaves fed with water (C) or D_5_-PAOx (D). Leaf segments of 8-day-old sorghum plants were incubated with either water or 30 μM of aldoximes for 24 h.

It was shown that nitrilase can take benzyl cyanide as a substrate in poplar, Arabidopsis, and sorghum (Jenrich *et al*., 2007; Urbancsok *et al*., 2018; Günther *et al*., 2018). Thus, one possible model of PAOx-derived PAA biosynthesis would be that PAOx is converted to benzyl cyanide, which can then be acted upon by nitrilases and ultimately converted to PAA. To test this model, we analyzed PAA content after benzyl cyanide feeding in sorghum and detected increased levels of PAA (**Fig 6A**). This suggests that benzyl cyanide can positively affect PAA accumulation, possibly by serving as a precursor of PAA or by activating PAA biosynthesis.

**Figure 6.**
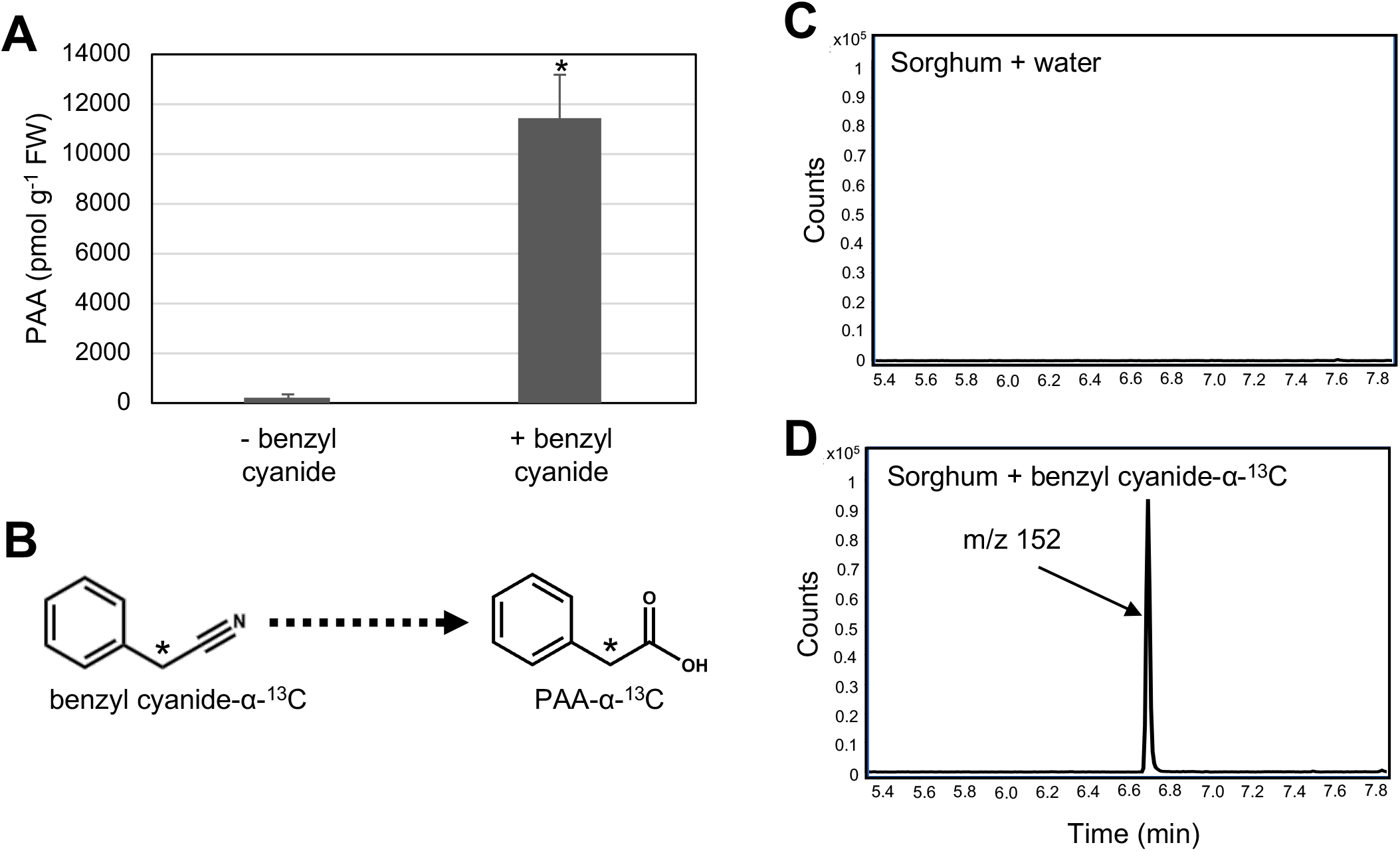
Sorghum can convert benzyl cyanide to PAA. (A) Free PAA concentration of sorghum leaves fed with water or benzyl cyanide (n=4). Bars represent mean, and error bars represent standard deviation. The means were compared by Student’s t-test and statistically significant differences (P < 0.05) are indicated by an asterisk (*). (B) Schematic of proposed conversion of benzyl cyanide-α-^13^C to PAA-α-^13^C. Location of the ^13^C atom in benzyl cyanide-α-^13^C and PAA-α-^13^C is denoted by an asterisk (*). (C-D) GC-MS chromatograms set for detection of the derivatized form of PAA-α-^13^C, methyl phenylacetate-α-^13^C (m/z 152) in sorghum leaves fed with water (C) or benzyl cyanide-α-^13^C (D). First leaves of 8-day-old sorghum plants were incubated with either water or 30 μM of benzyl cyanide for 24 h.

To determine if benzyl cyanide is converted to PAA, we analyzed sorghum plants after feeding with labeled benzyl cyanide-α-^13^C. To quantify and detect PAA using GC-MS, PAA must be derivatized with trimethylsilyldiazomethane to yield the volatile compound methyl phenylacetate. This derivatization results in an increase of 14 atomic mass units. Hence, for GC-MS detection and quantification of unlabeled and labeled PAA, mass spectrometers were set to detect compounds with the adjusted m/z values. A compound with m/z 152 was detected after feeding with benzyl cyanide-α-^13^C but not water (**Fig 6C-D**). This mass-to-charge ratio matched that of methyl phenylacetate-α-^13^C, the derivatized form of PAA-α-^13^C (**Fig 6B**), which is expected if benzyl cyanide acts as a precursor of PAA.

To determine if these results and pathways are shared between sorghum and maize, we also performed a similar series of feeding assays in maize. As with sorghum, feeding of maize with D_5_-PAOx but not water resulted in the accumulation of a compound with a mass-to-charge ratio and retention time identical to that of D_5_-benzyl cyanide (**Fig 7A-B**). Feeding with benzyl cyanide resulted in the accumulation of PAA in maize (**Fig 7C**), while feeding with benzyl cyanide-α-^13^C but not water resulted in the detection of a metabolite with a mass-to-charge ratio identical to that of methyl phenylacetate-α-^13^C (**Fig 7D-E**). Taken together, these results demonstrate that benzyl cyanide is a derivative of PAOx metabolism and serves as a direct precursor of PAA in both sorghum and maize (**Fig 8**).

**Figure 7.**
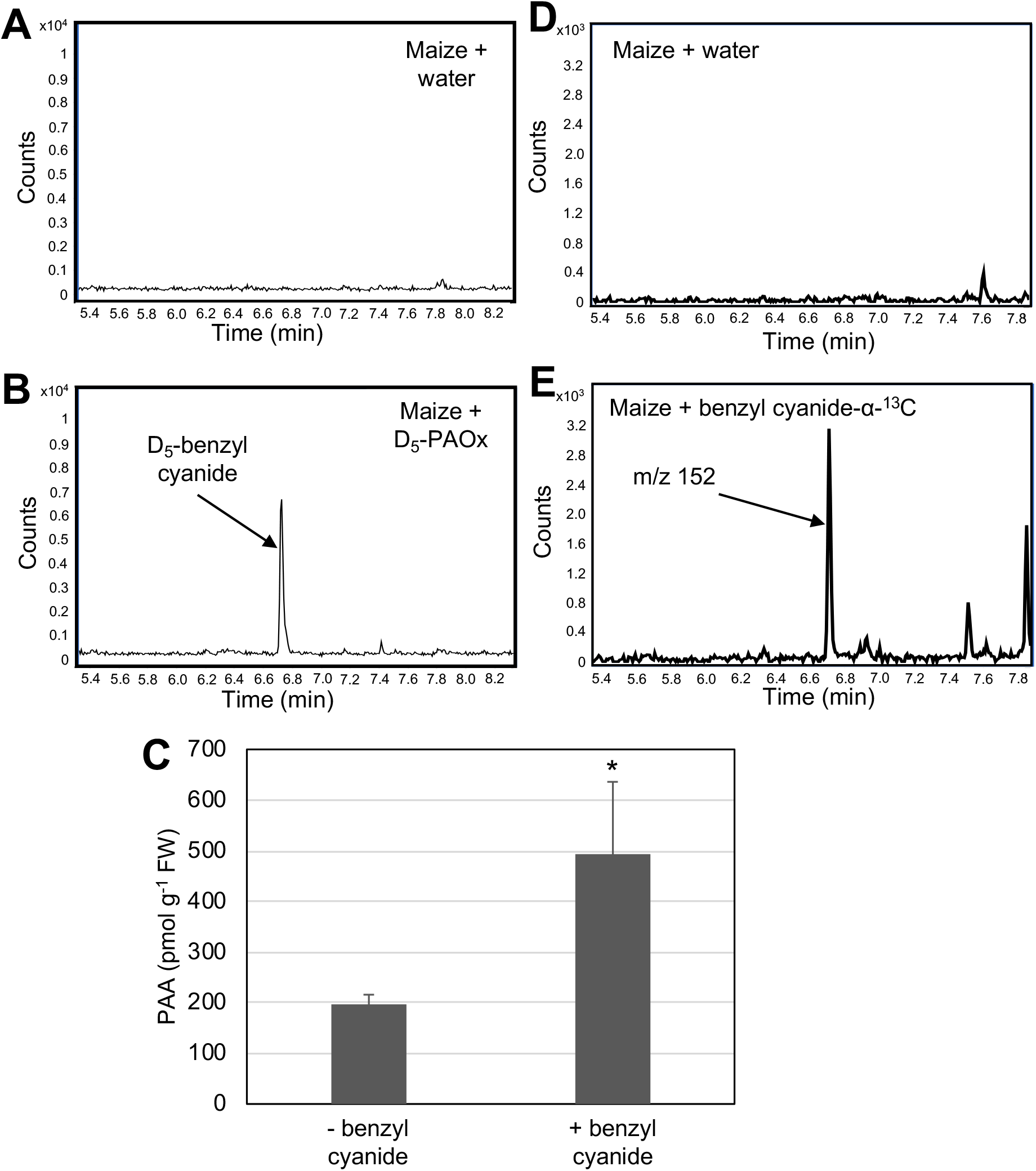
Production of benzyl cyanide and PAA in maize fed with PAOx. (A-B) GC-MS chromatograms set for detection of D_5_-benzyl cyanide (m/z 123) in maize leaves fed with water (A) or D_5_-PAOx (B). (C) Free PAA content in water-versus benzyl cyanide-fed 8-day-old maize plants (n=3). Bars represent mean, and error bars represent standard deviation. The means were compared by Student’s t-test and statistically significant differences (P<0.05) are indicated by an asterisk (*). (D-E) GC-MS chromatograms set for detection of the derivatized form of PAA-α-^13^C, methyl phenylacetate-α-^13^C (m/z 152) in maize leaves fed with water (D) or benzyl cyanide-α-^13^C (E). First leaves of 8-day-old maize plants were incubated with either water or 30 μM of aldoxime or benzyl cyanide for 24 h.

**Figure 8.**
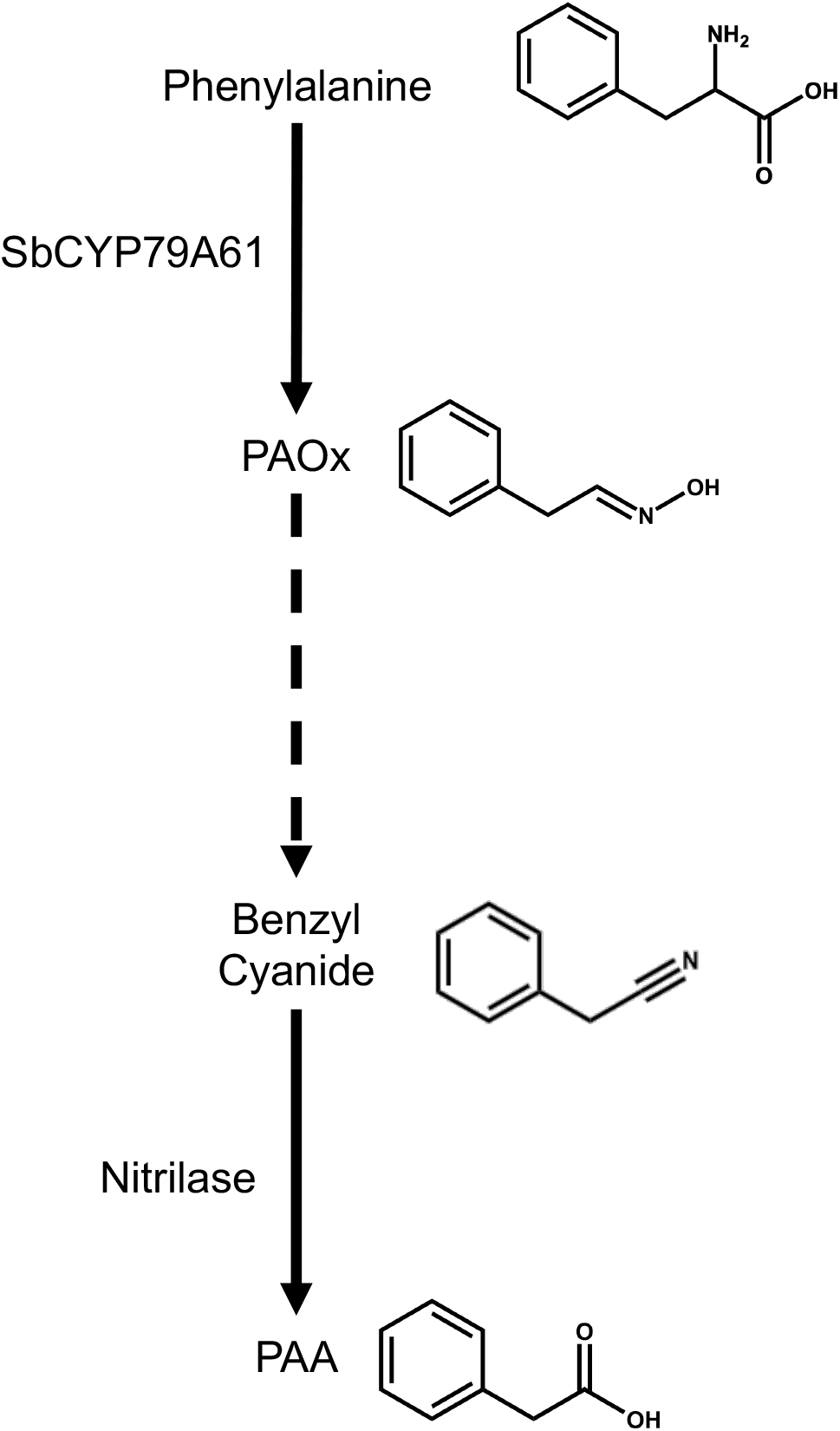
Proposed model of PAOx metabolism in sorghum. In sorghum, PAOx production from phenylalanine is catalyzed by SbCYP79A61. Then, through a yet uncharacterized rection(s), PAOx is converted to benzyl cyanide. Benzyl cyanide is then in turn converted to PAA. Even though the reactions linking benzyl cyanide to PAA are still unknown, it is likely that nitrilase is involved in this process. This pathway of PAOx metabolism is likely shared between sorghum and maize. Note that this work and model do not exclude the possibility of additional derivates of PAOx metabolism being present in sorghum and maize, such as cyanogenic glycosides.

## DISCUSSION

Here we identified SbCYP79A61 as an enzyme capable of generating PAOx in sorghum. Despite high amino acid sequence similarity between ZmCYP79A61 and SbCYP79A61 (**Fig S1**), the two enzymes had differing substrate specificities, with SbCYP79A61 having activity towards phenylalanine but little or no activity towards tryptophan, whereas ZmCYP79A61 produces both IAOx and PAOx (**Fig 2, 3**). An additional point of interest with the monocot CYP79s is their clustering into two distinct clades (**Fig 1**). One clade contains the aromatic-aldoxime-producing enzymes SbCYP79A1, SbCYP79A61 and ZmCYP79A61 and is part of a larger clade that contains aromatic-aldoxime-producing enzymes from monocots and dicots. The second, larger clade consists of the remaining three maize and seven sorghum CYP79 enzymes. Overall, these CYP79s except for SbCYP79A95 have low expression compared to their characterized counterparts (**Fig S2**) (Irmisch *et al*., 2015) and the enzymes form a clade separate from the examined CYP79s capable of producing aromatic or aliphatic aldoximes (**Fig 1**), which makes prediction of their activity difficult. Further study is required to identify their catalytic activities.

Besides CYP79 enzymes, other FMOs have been found to catalyze aldoxime production. Specifically, Thodberg *et al*. (2020) identified a fern oxime synthase (FOS1) within the species *Phlebodium aureum* and *Pteridium aquilinum* capable of catalyzing either one or two *N*-hydroxylations on phenylalanine to generate *N*-hydroxyphenylalanine and PAOx, respectively. This study identified the first instance of a non-CYP79 enzyme to catalyze amino acid to oxime conversion and provided a mechanism by which cyanogenic ferns that do not possess CYP79 enzymes can generate cyanogenic glycosides (Harper *et al*., 1976; Santos *et al*., 2005). The existence of FOS1 greatly expands the number of species potentially able to generate aldoximes, and the recent discovery of a new clade of FMOs capable of catalyzing *N*-hydroxylations (Liscombe *et al*., 2022) provides an additional target for identifying aldoxime production enzymes in both lower and higher plants, including sorghum.

While analyzing the potential role of SbCYP79A61 in sorghum, we determined that PAOx-derived PAA biosynthesis occurs in sorghum (**Fig 4**) and identified benzyl cyanide as an intermediate of PAOx-derived PAA in sorghum and maize (**Fig 5-8**). It has been proposed that nitriles such as indole-3-acetonitrile (IAN) are intermediates of aldoxime-derived auxin biosynthesis, based on the observation that Arabidopsis fed with labeled IAOx produces labeled IAN (Sugawara *et al*., 2009). Given that nitriles such as IAN and benzyl cyanide are products of glucosinolate degradation, it was unclear if IAN or benzyl cyanide is indeed an intermediate of aldoxime-derived auxin biosynthesis bypassing glucosinolate catabolism in Arabidopsis. However, our study revealed that benzyl cyanide is directly converted from PAOx and an intermediate of the PAOx-derived PAA biosynthesis in monocots. As both PAOx and benzyl cyanide production occurs widely in the plant kingdom and are induced by stressors (Irmisch *et al*., 2013; McCormick *et al*., 2014; Luck *et al*., 2016; Günther *et al*., 2018; Dhandapani *et al*., 2019; Liao *et al*., 2020), it is likely that PAOx-derived PAA biosynthesis contributes significantly to the pool of PAA in plants under stress. CYP71 enzymes have been shown to convert aldoximes to nitriles in various species (Nafisi *et al*., 2007; Klein *et al*., 2013; Yamaguchi *et al*., 2014, 2016; Irmisch *et al*., 2014; Liao *et al*., 2020). However, further study is needed to identify enzymes responsible for the conversion of PAOx to benzyl cyanide in monocots and determine if IAN is an intermediate of the IAOx-derived IAA biosynthesis in plants.

We detected a significant difference in PAA and benzyl cyanide accumulation upon PAOx feeding in sorghum, with the PAA concentration increasing to about 2,000 pmol/g fresh weight (**Fig 4A**), while benzyl cyanide concentration remained relatively low at less than 120 pmol/g fresh weight (**Fig 5A**). On the other hand, the magnitude of PAA accumulation upon benzyl cyanide feeding in sorghum and maize were significantly different, with PAA concentration increasing over 40-fold in sorghum (**Fig 6A**), but only around 2.5-fold in maize (**Fig 7C**). Even though we used the same age of leaf segments for our feeding experiment, maize and sorghum develop differently, and this difference could affect the feeding efficiency. It is also possible that differences in the activities of maize and sorghum nitrilases contribute to these differences. Maize and sorghum have two and three different nitrilase enzymes, respectively (Park *et al*., 2003; Jenrich *et al*., 2007) and previous studies have shown that these enzymes can form heterodimers with altered substrate affinities (Jenrich *et al*., 2007; Kriechbaumer *et al*., 2007). Both sorghum SbNIT4A and the SbNIT4A/SbNIT4B2 heterodimer have been shown to have activity towards benzyl cyanide, with the heterodimer having a K_m_ value of 0.19 mM and a V_max_ of 908 nkat (mg protein)^-1^ (Jenrich *et al*., 2007). The affinity of sorghum nitrilases for benzyl cyanide is equivalent to or higher than that of the Arabidopsis nitrilases (NIT1, NIT2, and NIT3), which have K_m_ values ranging between 0.14-1 mM and V_max_ values ranging from 40-1400 pkat (mg protein)^-1^ (Vorwerk *et al*., 2001). Initial characterization of the maize nitrilases ZmNIT1 and ZmNIT2 demonstrated that they have activity towards IAN (Park *et al*., 2003). Mukherjee *et al*. (2006) later showed that ZmNIT2 could hydrolyze benzyl cyanide into its corresponding acid and amide, although their experimental set up did not allow for quantification of substrate specificity. It is possible that the sorghum nitrilases have high affinity for benzyl cyanide, causing flux towards PAA to be relatively high and resulting in only low levels of benzyl cyanide accumulation and efficient conversion of benzyl cyanide to PAA in sorghum. Further research is needed to analyze the role nitrilase activity plays in PAOx-derived PAA biosynthesis.

In addition to PAA, PAOx can act as a precursor for cyanogenic glycosides. The PAOx-derived cyanogenic glycosides, amygdalin and prunasin, occur naturally in many species such as cassava, apple, and several *Prunus* species (Berenguer-Navarro *et al*., 2002; Yamaguchi *et al*., 2014; Bolarinwa *et al*., 2015, 2016; Thodberg *et al*., 2018). Dhurrin is the major cyanogenic glycoside in sorghum, but Bolarinwa *et al*. (2016) reported that sorghum grains also contain amygdalin. As other studies have not corroborated this observation (Pičmanová *et al*., 2015; Thodberg *et al*., 2018; Jaszczak-Wilke *et al*., 2021), amygdalin production may be genotype-specific or stress induced. Further study is needed to identify other aldoxime derivatives in monocots.

In summary, analysis of the sorghum genome for *CYP79* homologs using phylogenetic analysis resulted in the identification of the enzyme SbCYP79A61 that catalyzes the formation of PAOx. Furthermore, data generated from feeding assays and mass spectrometric analyses of sorghum and maize plants demonstrated that sorghum is capable of PAOx-derived PAA biosynthesis, and that benzyl cyanide serves as an intermediate of this pathway in both sorghum and maize. These results suggest that PAOx and PAOx-derived metabolism may play a role in fine-tuning plant responses to various stressors.

## Abbreviations

PAA: phenylacetic acid
PAOx: phenylacetaldoxime

## Supporting Information

Table S1. List of amino acid sequence identities of SbCYP79A61 with other CYP79 enzymes.

Figure S1. Alignment of SbCYP79A61 protein sequences to maize ZmCYP79A61.

Figure S2. Tissue-specific expression profile of *CYP79* genes in *Sorghum bicolor*.

## Data Availability

All data supporting the findings of this study are available within the paper and within its supplementary materials published online.

## Acknowledgements

This work was supported by the United States Department of Agriculture (USDA)-National Institute of Food and Agriculture Hatch (005681) and a startup fund from the Horticultural Sciences Department and Institute of Food and Agricultural Sciences at the University of Florida to JK. The United States Department of Agriculture (USDA)-Agricultural Research Service Project number 6036-11210-001-00D and the USDA-National Institute of Food and Agriculture-Specialty Crop Research Initiative grant 2018-51181-28419 to AB.

## Author Contributions

R.D, V.C.P, W.V., A.G., A.K.B., J.K. designed the research project; R.D, V.C.P, A.B, B.T., E.S.A.W., J.M. performed the experiments and analyzed the data; V.C.P and J.K. wrote the manuscript. All authors read and agreed with the manuscript.

